# NanoTox: Development of a parsimonious *in silico* model for toxicity assessment of metal-oxide nanoparticles using physicochemical features

**DOI:** 10.1101/2021.02.22.432301

**Authors:** Nilesh AnanthaSubramanian, Ashok Palaniappan

## Abstract

Metal-oxide nanoparticles find widespread applications in mundane life today, and cost-effective evaluation of their cytotoxicity and ecotoxicity is essential for sustainable progress. Machine learning models use existing experimental data, and learn the relationship of various features to nanoparticle cytotoxicity to generate predictive models. In this work, we adopted a principled approach to this problem by formulating a feature space based on intrinsic and extrinsic physico-chemical properties, but exclusive of any *in vitro* characteristics such as cell line, cell type, and assay method. A minimal set of features was developed by applying variance inflation analysis to the correlation structure of the feature space. Using a balanced dataset, a mapping was then obtained from the normalized feature space to the toxicity class using various hyperparameter-tuned machine learning models. Evaluation on an unseen test set yielded > 96% balanced accuracy for both the random forest model, and neural network with one hidden layer model. The obtained cytotoxicity models are parsimonious, with intelligible inputs, and include an applicability check. Interpretability investigations of the models yielded the key predictor variables of metal-oxide nanoparticle cytotoxicity. Our models could be applied on new, untested oxides, using a majority-voting ensemble classifier, NanoTox, that incorporates the neural network, random forest, support vector machine, and logistic regression models. NanoTox is the very first predictive nanotoxicology pipeline made freely available under the GNU General Public License (https://github.com/NanoTox).

## INTRODUCTION

Nanotechnology has delivered the promise of ‘plenty of room at the bottom’ with transformative applications for human welfare [1]. The distinctive properties of nanoscale materials have been indispensable in industrial and medical applications, including the delivery of biologically active molecules, and development of biosensors for human health and disease [2]. Engineered metal-oxide nanoparticles are characterized by a concentration of sharp edges, and lend themselves to a variety of uses (for e.g, [3]). However there is a potential caveat to nanobiotechnology: the differential nanoscale behaviour of nanomaterials also obtains emergent toxic side-effects in the biological domain and ecological realm [4–7]. These hazards are related to the capacity of nanomaterials to engender free radicals in the cellular milieu, which inflict damaging oxidative stress. Such events could trigger inflammatory responses, which could balloon out of control leading to apoptosis and cytotoxicity [8–11], and genotoxicity [12].

The mundane use of nanoparticles has necessitated vigorous safety assessment of toxicity, in the interests of sustainable progress [13–16]. Such methods could also help discern safe-by-design principles that could guide adjustments to the nanoparticle formulation, and thereby mitigate adverse effects at the source. Intelligent and alternative testing strategies could accelerate rational design of nanoparticles for optimal functionality and minimal toxicity [17–20]. Various computational methods have been applied to predicting toxicity of engineered nanomaterials [21–30], but with the accumulation of high-quality data, machine learning methods have shown the most promise [31]. Such techniques provide a non-invasive ‘instantaneous’ readout of nanoparticle toxicity [32–34], and originate from the evolution of QSAR models [35]. Machine learning models of nanoparticle toxicity have tended to be either generalized [36] or tissue-specific [37,38], and are built from experimental toxicity data that have been scored, standardised and curated into databases like the Safe and Sustainable Nanotechnology db (S2NANO) [39–41].

Earlier studies have tended to neglect systematic multicollinearity among the predictor variables, which would lead to confounding and data snooping. Secondly, gross imbalance between the numbers of nontoxic and toxic instances usually exists, which could lead to overfitting to the ‘nontoxic’ class [42]. Third, we were motivated to develop a model that would be agnostic of *in vitro* characteristics, such as cell line, cell type, and assay method. A truly general model of nanoparticle cytotoxicity, independent of *in vitro* factors, would lead to significantly broader interpretability and wider applicability [43]. Our study departs also from the notion that tissue-specific models are superior to generalized models [38], and demonstrates that model interpretability is best achieved using a minimal non-redundant feature space, consistent with Occam’s parsimony. Furthre, with a view to increasing reliability, we have deployed the insights from our study into a majority-voting ensemble classifier. Finally, the end-to-end pipeline of our work, including the ensemble classifier, is made freely available as a user-friendly open-source nanosafety prediction system, NanoTox, under GNU GPL (https://github.com/NanoTox). All implementations were carried out in R (www.r-project.org).

## METHODS

### Problem and dataset

*In vitro* parameters such as cell-type, cell line, cell origin, cell species, and type of assay, could be extraneous to modelling the intrinsic hazard posed by a nanoparticle to cellular viability and the environment. This motivated us to formulate the problem in a feature space devoid of biological predictors. The machine learning task is stated as: given a certain nanoparticle at a certain dose for a certain duration, would its administration prove cytotoxic? To address this problem, we used a hybrid dataset building on the physico-chemical descriptors and toxicity data found in Choi *et al*’s study [36]. All *in vitro* features were removed from the dataset, as noted above. Extrinsic physico-chemical properties, namely dosage and exposure duration were retained [44]. The periodic table properties of metal-oxide nanoparticles published in Kar et al [45] were used to augment the dataset. Only complete cases were considered in the process of matching the two datasets. This process yielded a final dataset of 19 features of five metal-oxide nanoparticles: Al_2_O_3_, CuO, Fe_2_O_3_, TiO_2_ and ZnO (Table 1). Cytotoxicity was used as the outcome variable, encoded as ‘1’ (true) if measured cell viability was < 50% with respect to the control, and ‘0’ (false) otherwise. The dataset is available on NanoTox.

**Table 1.**
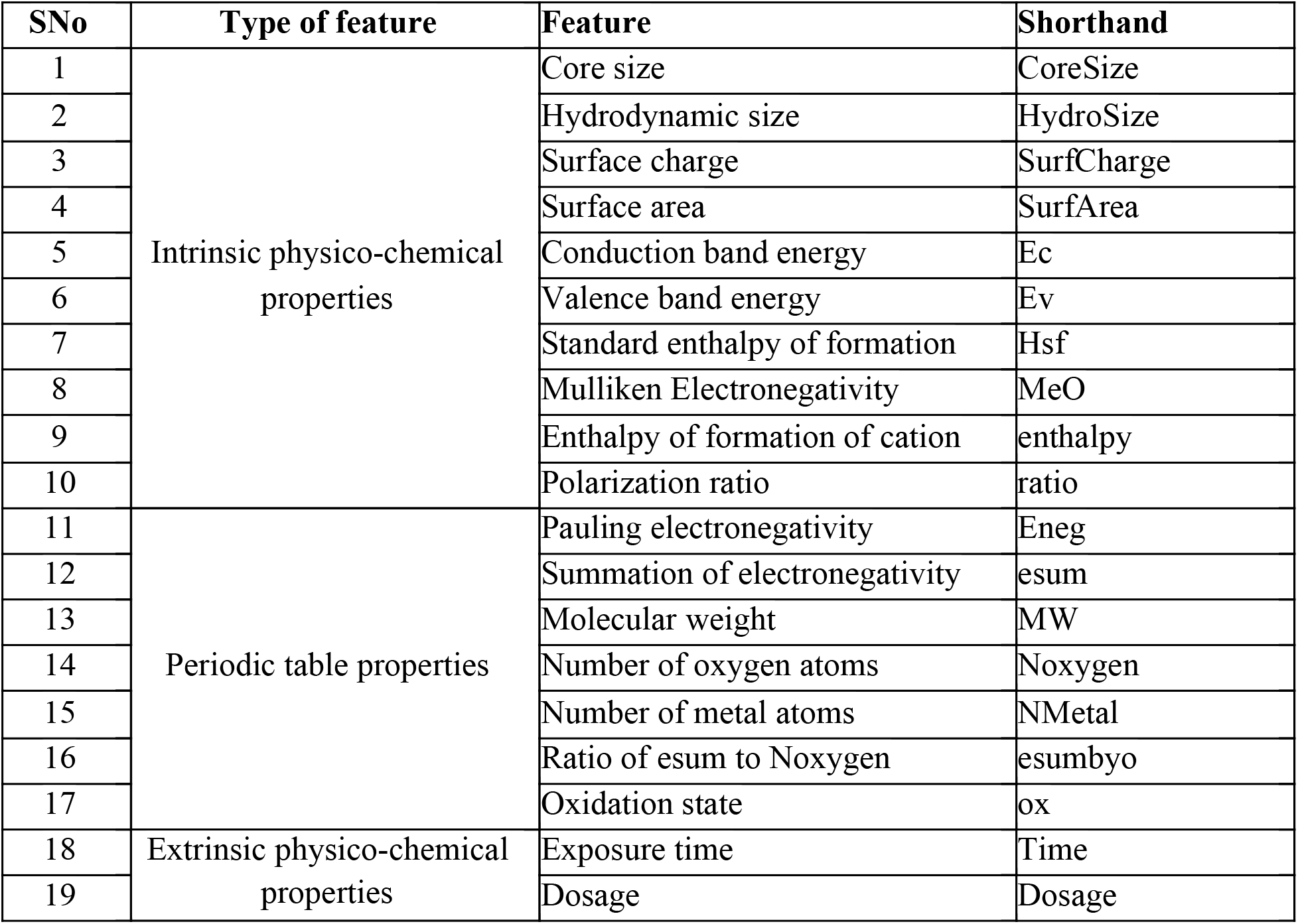
The 19 physico-chemical features of MeOx nanoparticles considered in our study.

| SNo | Type of feature | Feature | Shorthand |
| --- | --- | --- | --- |
| 1 | Intrinsic physico-chemical properties | Core size | CoreSize |
| 2 |  | Hydrodynamic size | HydroSize |
| 3 |  | Surface charge | SurfCharge |
| 4 |  | Surface area | SurfArea |
| 5 |  | Conduction band energy | Ec |
| 6 |  | Valence band energy | Ev |
| 7 |  | Standard enthalpy of formation | Hsf |
| 8 |  | Mulliken Electronegativity | MeO |
| 9 |  | Enthalpy of formation of cation | enthalpy |
| 10 |  | Polarization ratio | ratio |
| 11 | Periodic table properties | Pauling electronegativity | Eneg |
| 12 |  | Summation of electronegativity | esum |
| 13 |  | Molecular weight | MW |
| 14 |  | Number of oxygen atoms | Noxygen |
| 15 |  | Number of metal atoms | NMetal |
| 16 |  | Ratio of esum to Noxygen | esumbyo |
| 17 |  | Oxidation state | ox |
| 18 | Extrinsic physico-chemical properties | Exposure time | Time |
| 19 |  | Dosage | Dosage |

### Elimination of multicollinearity

A simple inspection of the properties in Table 1 suggested the existence of correlated features. Correlated features would adversely impact model performance as well as complicate model interpretation. Multicollinearity is an even deeper problem in the pursuit of a non-redundant feature space [46]. The existence of highly correlated (Pearson’s rho >= 0.9) variables was first ascertained. To address multicollinearity, we used a systematic variance inflation factor (vif) analysis. Each independent variable was regressed on all the other independent variables in turn, and the goodness-of-fit of the models (fraction of variance explained; R^2^) were estimated. The vif-score for each independent variable was then calculated using eqn. (1). In each iteration of the vif analysis, the variable in the current set that had the largest vif score when regressed on all the other variables was eliminated. This process was continued until a set of variables all of whose vif scores < 5.0 was obtained. Note that a vif score of 1.0 is possible only when a variable is perfectly independent of all other variables (all pairwise Pearson’s rho identically zero).

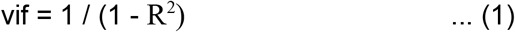

### Feature transformation

The feature space could be vulnerable to heteroscedastic effects, given the varying scales for the variables. It is necessary to pre-process and prevent features with large variances from swamping out the rest. Positively skewed features could be stabilized using the log transformation. Ec values, which are negative, were first offset by +6.17, then log-transformed. Dosage spanned many orders-of-ten, and was log_10_-transformed. Exposure time spanned orders-of-two magnitude, so we performed a log_2_ transformation. Surface charge whose values could be either positive or negative, was standardized (i.e., Z-transformed). All the other features were log-transformed (to the base *e*).

### Class rebalancing

The cost of missing a toxic instance is manifold higher than the cost of missing a non-toxic instance, and the imbalance between toxic vs nontoxic instances could exacerbate this problem. In such situations, where the essential problem is to learn the minority outcome class effectively, resampling techniques could be useful [47]. We addressed the class skew problem using Synthetic Minority Over-Sampling TEchnique (SMOTE) [48]. SMOTE synthesises new minority samples from the existing ones, without influencing the instances of the majority class, thereby increasing the number of “toxic” instances relative to the number of “non-toxic” instances. Balancing the dataset thus would normalise the learning bias arising from unequal representation of the outcome classes.

### Predictive Modelling

The overall workflow of our approach is summarized in Figure 1. The normalized, balanced dataset was randomly split into an 70:30 train:test ratio stratified on the outcome variable [49]. Towards this, a variety of classification algorithms were tried and tested, namely logistic regression [50], random forests [51], SVMs [52], and neural networks [53,54]. Table 2 shows these details and the classifier-wise hyperparameters considered in our work. The optimal values of the hyperparameters were found using 10-fold internal cross-validation [55]. The performance of each optimized model was evaluated on the unseen test set. In order to penalize false positives and false negatives equally, we used an objective measure of performance:

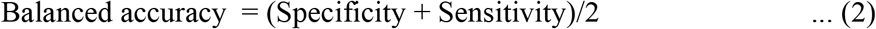

**Figure 1.**
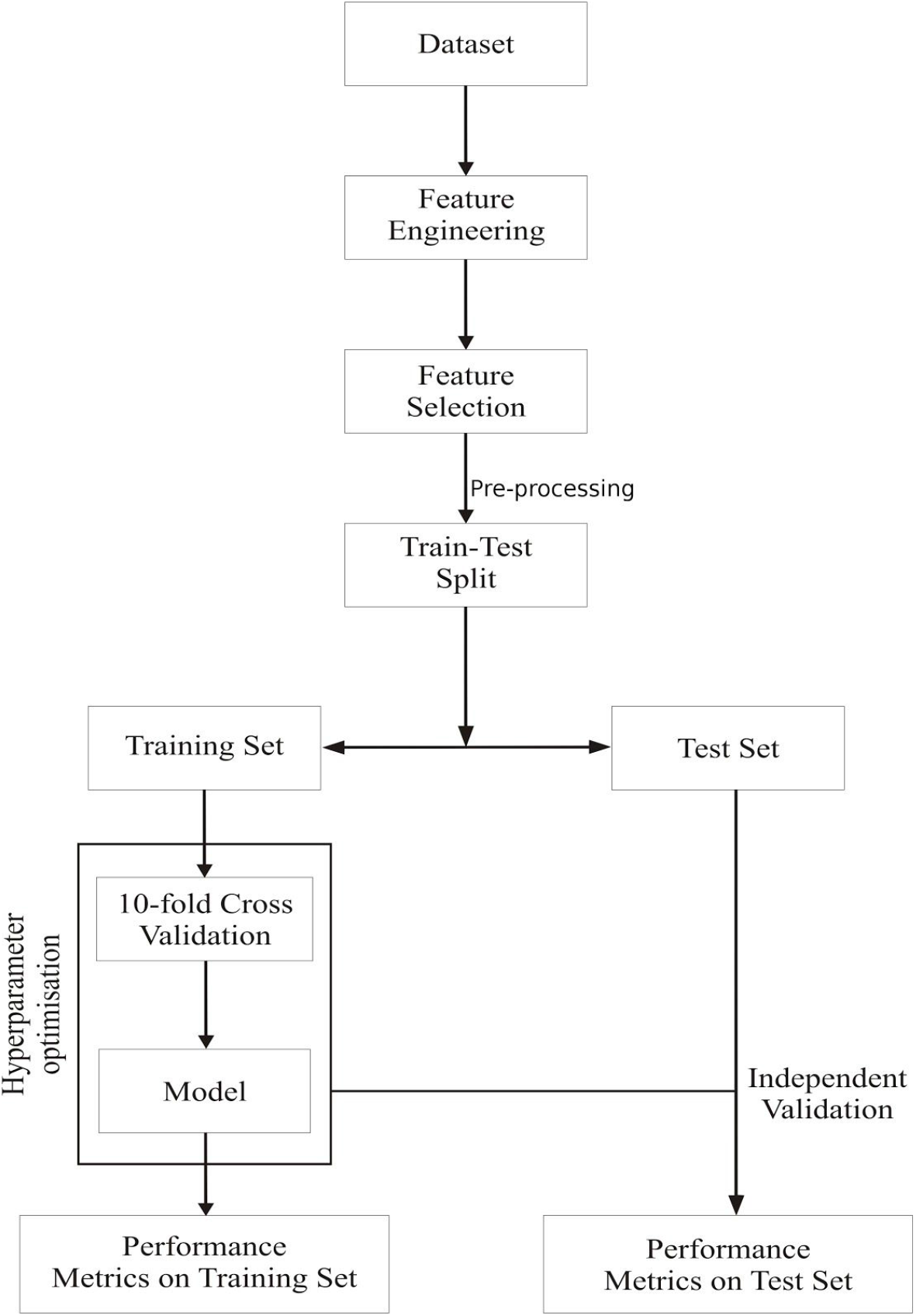
The workflow of the study upto predictive modelling. Pre-processing refers to both normalization and class balancing.

**Table 2.** Classifiers used in our study and their respective hyperparameters. m_try_ represents the number of features used for each split in the random forest model.

| No. | Classifier | Type / Basis | Package / function | Hyperparameters | Optimisation |
| --- | --- | --- | --- | --- | --- |
| 1 | Logistic regression | Algebraic | glm | Threshold (= 0.5) | n/a |
| 2 | Random forest | Rule-based | randomForest | 1. #trees (= 500)<br>2. $m_{\text{try}}$ | caret::train |
| 3 | Support vector machine | Geometric | e1071 | 1. Kernels (linear, radial, polynomial)<br>2. Cost<br>3. Gamma<br>4. Degree | e1071::tune |
| 4 | Neural networks | Connectionist | RSNNS | 1. #hidden layers = 1,2<br>2. Size of each hidden layer<br>3. Decay rate | caret::train, caret::mlpML |

### Applicability domain

The specification of the applicability boundaries of machine learning models would increase their reliability and utility [43]. This would define the perimeter of model generalization to new instances, and safeguard against application to atypical data. We used a Euclidean nearest-neighbor approach to define the applicability domain (AD) of the machine learning models [56]. For each instance in the training set, its distances to all the other training instances were found. The nearest neighbours of each instance are then defined as the k smallest values from this set, where k is an integer parameter set to the square-root of the number of instances in the training set. The mean distance of an instance to its k-nearest neighbours is found, and this process is repeated for all instances to yield the sampling distribution of these mean distances. The mean and standard deviation of this sampling distribution were designated as μ_k_ and σ_k_, respectively. The applicability domain is then defined as follows:

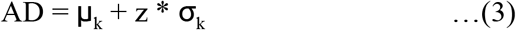

where z is an empirical parameter (related to the z-distribution) that characterises the width of belief in the model, and here set to 1.96.

## RESULTS

Our dataset consisted of 483 instances of the five metal-oxide nanoparticles with 19 features and one outcome variable. Correlogram plots identified the existence of high correlation among these 19 variables (Figure 2; see Figure S1 for a zoomed correlogram of the periodic table properties). Three clusters of high correlation were revealed: one cluster of enthalpy, Hsf, ratio, ox, Noxygen, and esumbyo; a second cluster of Ec and Ev; and a third cluster of esum, NMetal and MW. Based on the vif analysis, we were able to obtain a feature space of just nine uncorrelated non-redundant variables (Table 3). The highest vif of any variable in this feature space was < 2.02, indicating little residual multicollinearity (Figure 3). This optimal feature space included two periodic table properties (Eneg, NOxygen), five other intrinsic physico-chemical properties (CoreSize, HydroSize, SurfArea, SurfCharge, Ec), and both the extrinsic physico-chemical properties (Dose, Time). This final dataset of 483 instances with nine features and one outcome variable is available at NanoTox.

**Figure 2.**
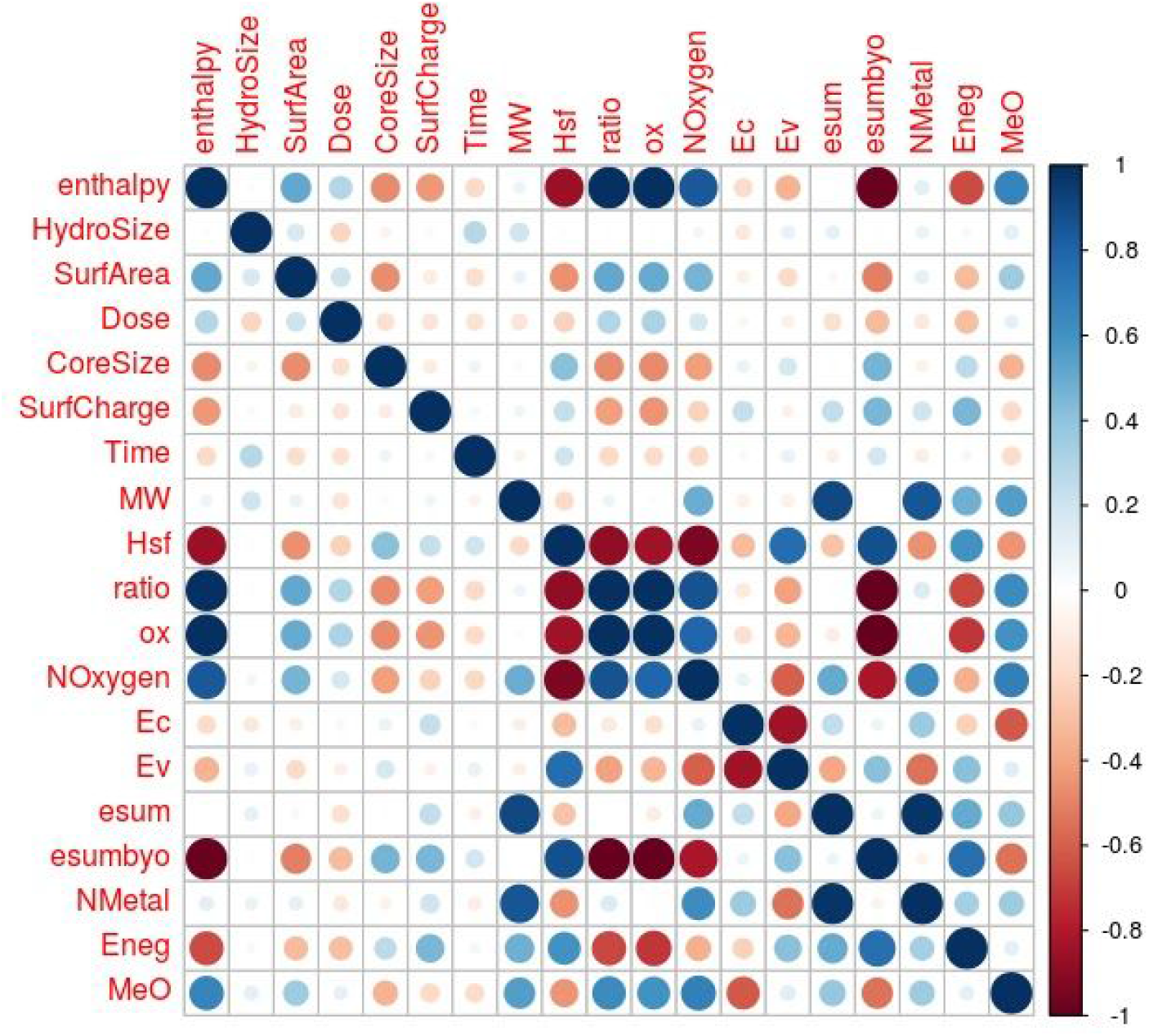
Correlogram of the 19 features. The correlation between a row feature and a column feature is shown by a dot in the corresponding cell. The size of the dot represents the magnitude of the correlation, and colour represents the sign of the correlation – blue: positive, and red: negative. White indicates a value near 0, i.e, independence.

**Figure 3.**
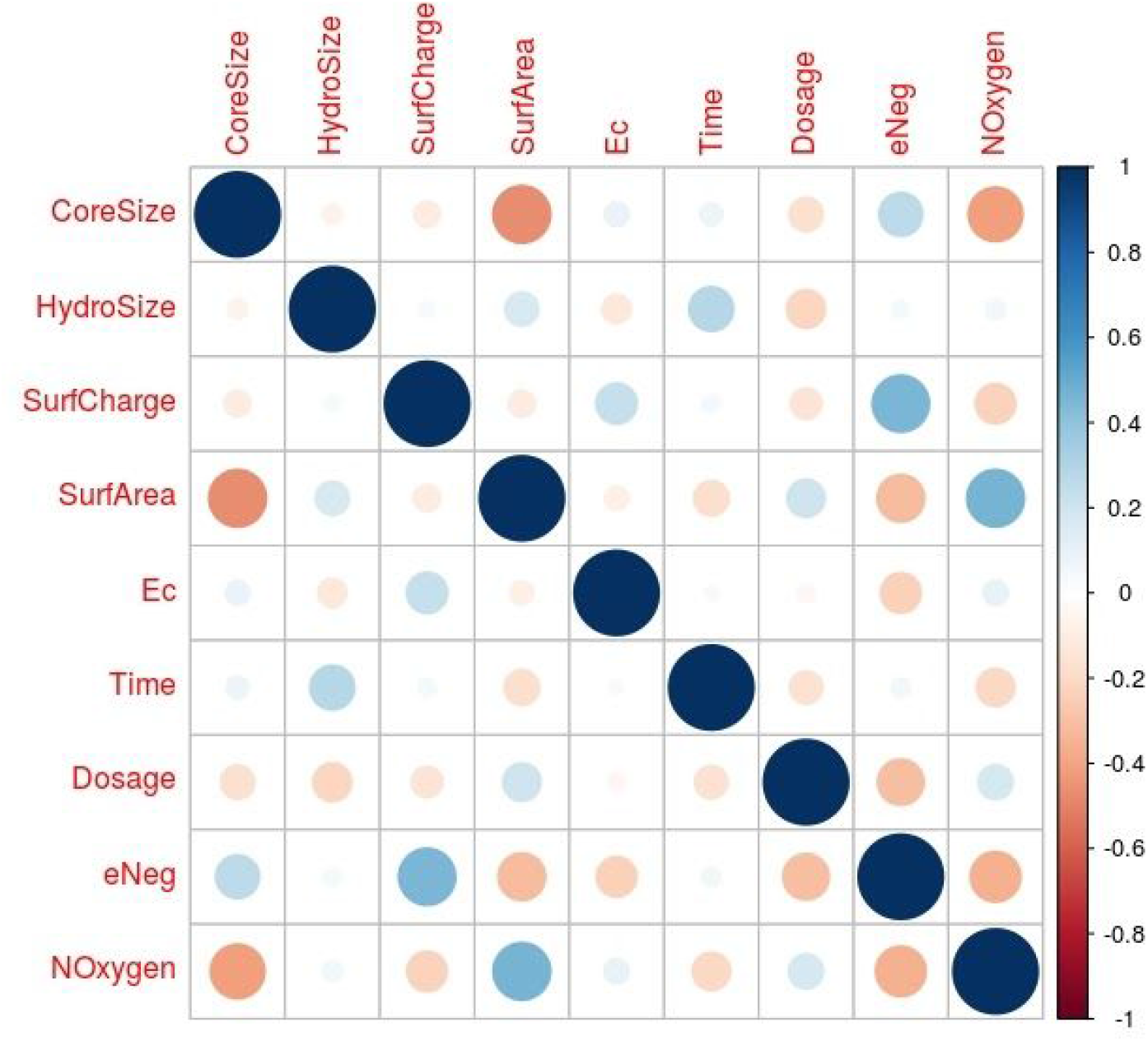
Correlogram of the optimised feature space. No subset of variables in this set is multicollinear.

**Table 3.** vif scores for the features in the final reduced set. The maximum vif score is ~ 2.0, corresponding to maximum R^2^ ~ 0.5 (cf. Eqn.(1)).

| S.No | Feature | Variance inflation factor |
| --- | --- | --- |
| 1 | CoreSize | 1.65 |
| 2 | HydroSize | 1.24 |
| 3 | SurfCharge | 1.85 |
| 4 | SurfArea | 1.58 |
| 5 | Ec | 1.50 |
| 6 | Time | 1.19 |
| 7 | Dose | 1.21 |
| 8 | Eneg | 2.02 |
| 9 | NOxygen | 1.60 |

The nine features were normalized, producing acceptable skew values for HydroSize, SurfArea, Ec, and Time (Table 4). The normalized dataset was partitioned using a random 70:30 split stratified on the outcome variable, providing a training dataset of 339 instances (with 55 ‘toxic’ instances), and an independent test dataset of 144 instances (with 23 ‘toxic’ instances). The training dataset (and not the test dataset) was balanced for the minority “toxic” instances using SMOTE resampling, yielding 165 ‘toxic’ and 220 ‘non-toxic’ instances, for a training dataset of 385 instances. This normalised and balanced dataset was used to train the various classifiers. The optimal hyperparameters of each classifier were determined using the R e1071 package for SVMs (Figure S2), and the R caret package for the neural networks, both one-layer (Figure 4) and two-layers (Figure S2). The full set of model-wise optimal hyperparameters could be found in Table S1. The trained, optimised classifiers were then evaluated on the unseen test dataset. All the models, except the SVM with polynomial kernel, achieved perfect sensitivity to the ‘toxic’ instances, i.e. all cytotoxic nanoparticles were classified correctly. The models were not perfectly specific to the non-toxic instances, however. On this basis, the random forest and neural network - one layer models outperformed all the others. They were each frustrated by eight false positives, yielding a balanced accuracy of 96.69%. All the classifiers achieved balanced accuracy > 90%. Table 5 summarises the performance of all the models on the test set. Five ‘non-toxic’ instances were classified incorrectly by all the models, representing refractory instances and constituting a challenge to perfect learning. One of these instances was only marginally viable (0.52), indicating the possible source of refractoriness.

**Figure 4.**
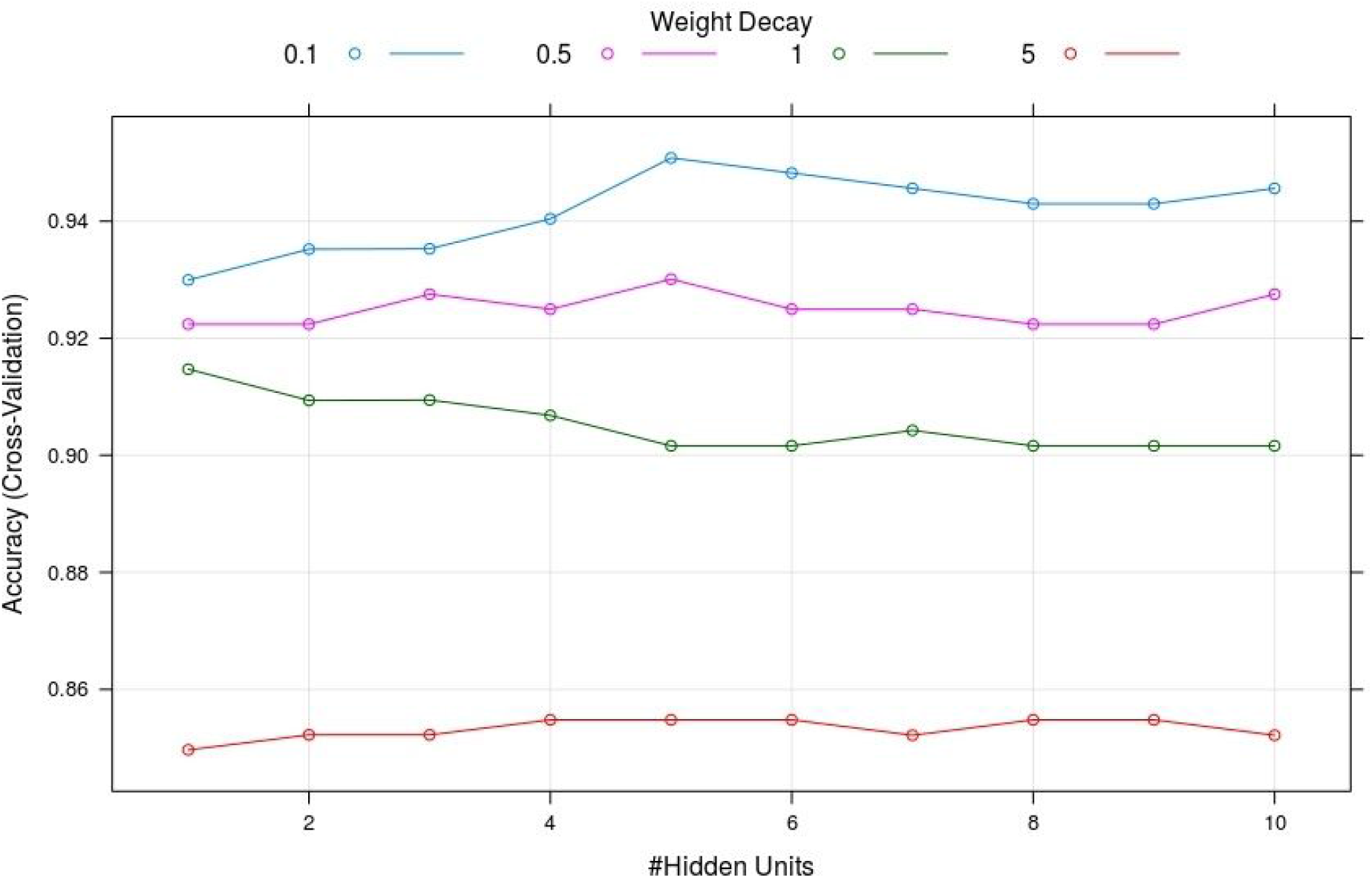
Hyperparameter tuning, for the neural network - 1 layer model. It is seen that the cross-validation accuracy is sensitive to the choice of the set of hyperparameters.

**Table 4.** Dataset normalization. Log-transformation was performed to the base *e*. Skewness was controlled, and the range of all predictors was brought into the same order of magnitude.

| Features | Skewness before | Type of Normalization | Skewness after | Range (min-max) |
| --- | --- | --- | --- | --- |
| CoreSize | 0.92 | Log | -0.2 | 2.01 - 4.82 |
| HydroSize | 1.76 | Log | 0.14 | 4.30 - 7.52 |
| SurfCharge | 0.45 | z-score | 0.45 | -1.62 - +1.98 |
| SurfArea | 2.14 | Log | -0.23 | 1.95 - 5.35 |
| Ec | 2.68 | Log, with offset | -0.23 | 0.00 - 1.54 |
| Time | 1.36 | Rescale, log | -0.48 | 0.00 - 4.58 |
| Dosage | 1.74 | Log10 | -1.5 | -5.00 - +2.48 |
| Eneg | 1.46 | Log | 1.26 | 0.43 - 0.64 |
| Noxygen | 0.66 | none | 0.66 | 1 - 3 |

**Table 5.** Performance of the various models. Models with balanced accuracy > 94% are highlighted.

| ID | Classifier | Train set |  |  | Test set |  |
| --- | --- | --- | --- | --- | --- | --- |
|  |  | Accuracy | Balanced accuracy | Cross-valid accuracy | Accuracy | Balanced accuracy |
| Model_1 | Logistic regression | 0.94 | 0.94 | 0.93 | 0.91 | **0.95** |
| Model_2 | Random forest | 0.98 | 0.98 | 0.94 | 0.94 | **0.97** |
| Model_3a | SVM-Linear | 0.94 | 0.95 | 1 | 0.9 | 0.94 |
| Model_3b | SVM-Radial | 0.94 | 0.94 | 1 | 0.86 | 0.92 |
| Model_3c | SVM-Poly | 0.98 | 0.98 | 1 | 0.84 | 0.85 |
| Model_4a | Neural network – 1L | 0.96 | 0.96 | 0.94 | 0.94 | **0.97** |
| Model_4b | Neural network – 2L | 0.96 | 0.95 | 0.95 | 0.91 | **0.95** |

### Deployment

The applicability domain was calculated with the normalised train data, prior to SMOTE balancing. Substituting k = 19 and z = 1.96 in eqn. (2) yielded the AD threshold = 2.23. About 95% of the test instances (i.e, 137 / 144 instances) were located within the AD radius. It must be noted that the misclassified instances did not coincide with these outliers. We have provided a workflow, deployment.R (available at NanoTox), for prediction on new, untested oxides. The prediction is executed by a majority-voting ensemble classifier [57], since bagging the predictions of the best models on the test set improved the performance to just five false positives (~ 98% balanced accuracy). Any new instance for classification supplied by the user is pre-processed (normalised), and its ‘typicality’ determined by calculating its distances to the instances in the original train data, and finding the mean, D_i_, of the 19 closest distances. If the D_i_, is greater than the AD threshold, then the instance is deemed atypical for requesting the ensemble model. Predictions are obtained using the top two models – the random forest and the neural network - one layer, and a consensus prediction is sought. In the absence of a consensus, an ensemble of the top five classifiers, all with balanced accuracy > 94% (highlighted in Table 5), is used. In the end, the majority prediction of the ensemble classifier is the predicted cytotoxicity of the given instance. Deployment.R automates this pipeline for a batch of new, untested oxides of any size. Furthermore, the RDS images of all the models trained in our study are provided on NanoTox, for the interested scientist.

## DISCUSSION

It is clear that SMOTE balancing made a difference in the ability of the classifiers to detect the under-represented ‘toxic’ instances. Filtering based on applicability domain and use of an ensemble classification strategy further mitigate model uncertainty given the ‘no free lunch’ theory. Benchmarking our results with Choi *et al*. [36], we see that the best model in each classifier from our work outperformed the corresponding best models of their work (Table 6). The overall best models in our work (random forest and neural network 1-layer) yielded a balanced accuracy of ~97% compared to 93% for their best overall model (‘neural networks’). All the five models from this work with balanced accuracy > 93% are deployed in an ensemble classifier to further mitigate uncertainty in prediction.

**Table 6.** Benchmarking. SVM (a), (b), and (c) correspond to linear, radial, and polynomial kernels. respectively. Neural networks (a) and (b) refer to one and two hidden layer(s), respectively. No information regarding model hyperparameters were available in Choi et al. [35]. The best-performing models from our work are highlighted.

| Model | Balanced accuracy (%) |  |
| --- | --- | --- |
|  | Choi et al. [35] | Present work |
| Logistic regression | 92 | 94.63 |
| Random forest | 91 | **96.69** |
| SVM | 91 | (a) 94.21 |
|  |  | (b) 91.74 |
|  |  | (c) 85.21 |
| Neural networks | 93 | (a) **96.69** |
|  |  | (b) 94.63 |

Figure 5 shows the best-performing neural network model, with connections weighted by their importance [58]. A variable importance plot of the random forest shows Dose as the most important predictor, followed by NOxygen, Eneg and Time (Figure 6(a)) [59]. A relative importance analysis of the neural network – 1-layer model obtains concurrence to these findings, and adds a direction to the favoured binary outcome (Figure 6(b)) [60,61]. Dose emerges as the key variable determining nanoparticle toxicity, and Time, HydroSize and Eneg are the other variables influencing the ‘toxic’ prediction. NOxygen emerges as the key predictor influencing the ‘nontoxic’ prediction, and SurfArea, Ec and CoreSize are the other predictors in this category. Logistic regression provides us with not just the effect size (coefficients) of the individual variables but also an estimate of their significance (in terms of the p-value of the coefficients) (Table S2). It is significant that the logistic regression model supports these findings, with few inconsistencies. Further the sign of the coefficient of each variable matches its direction of influence presented in Figures S3, S4. It is noteworthy that the two ‘periodic table’ properties (Eneg, Noxygen), and the quantum chemical property, Ec, show large effect sizes (coefficients) but poor significance, when all the other variables remain extremely significant.

**Figure 5.**
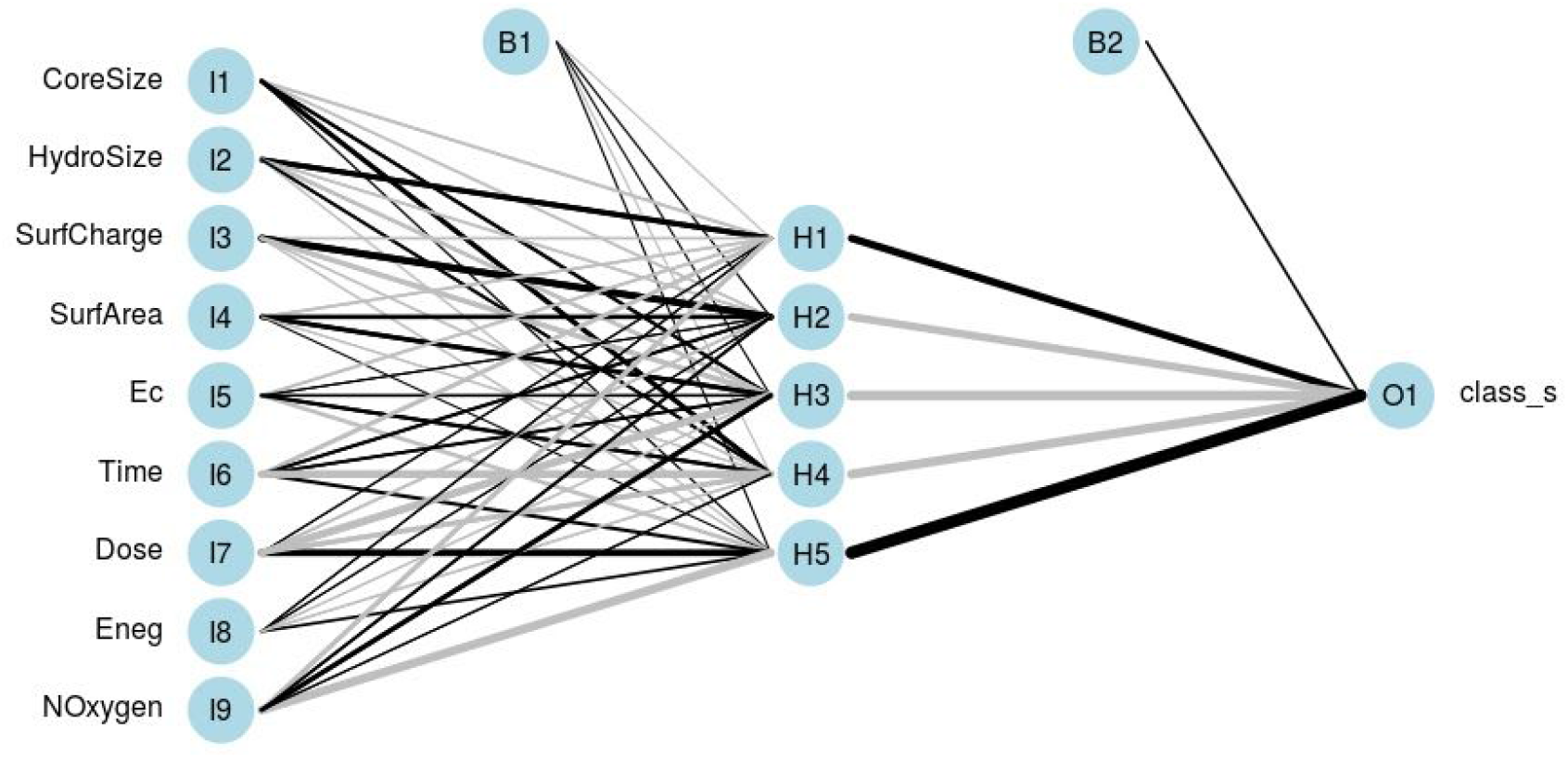
A schematic of the trained neural network-1 layer model, with the weights of the connections indicated by the linewidth. Black lines indicate positive weights, and gray lines indicate negative weights. Two bias units are seen, one for the hidden layer, and one for the output layer.

**Figure 6.**
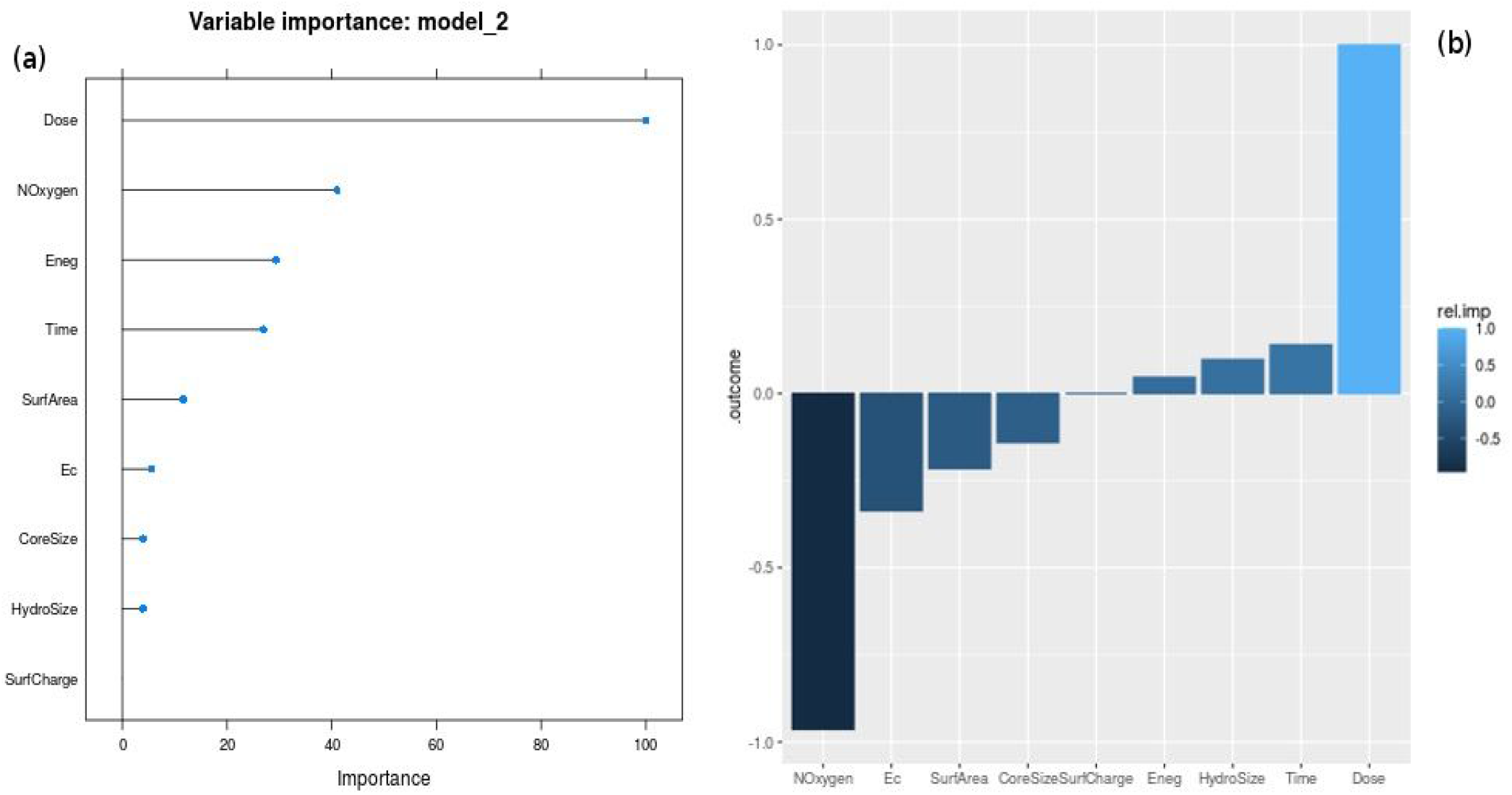
(a) Normalized variable importance for the Random Forest model computed with caret. Dose is by and far the attribute with the greatest effect on the toxicity in the Random Forest model. (b) Relative importance plot for the NeuralNet-1L. Positive values correspond to the ‘true’ (i.e, ‘toxic’) class, and negative values correspond to the ‘non-toxic’ class. It is seen that Dose and NOxygen exert the maximum importance on the outcome class, though in opposite directions.

Consensus among the models is necessary for explainable AI [62], and in this direction, we performed a Lek sensitivity analysis with the neural network 1-layer model [63]. How does the response variable change with changes in a given explanatory variable, given the context of the other explanatory variables? In investigating the effect of one explanatory variable, all the other explanatory variables are clustered into a specified number of lakes with like characteristics. While the unevaluated explanatory variables are held constant at the centroid of one lake cluster, the explanatory variable of interest is sequenced from minimum to maximum in 100 quantile steps, with the response variable predicted at each step, yielding a sensitivity curve. This process is iterated for each lake of the unevaluated explanatory variables, yielding the sensitivity profile of the response variable with respect to the specific explanatory variable in the context of the unevaluated explanatory variables. We set the number of clusters to ten, to visualize a sufficient number of the response curves for each explanatory variable. In this way, the sensitivity profiles of the response variable are obtained for each predictor (Figure 7).

**Figure 7.**
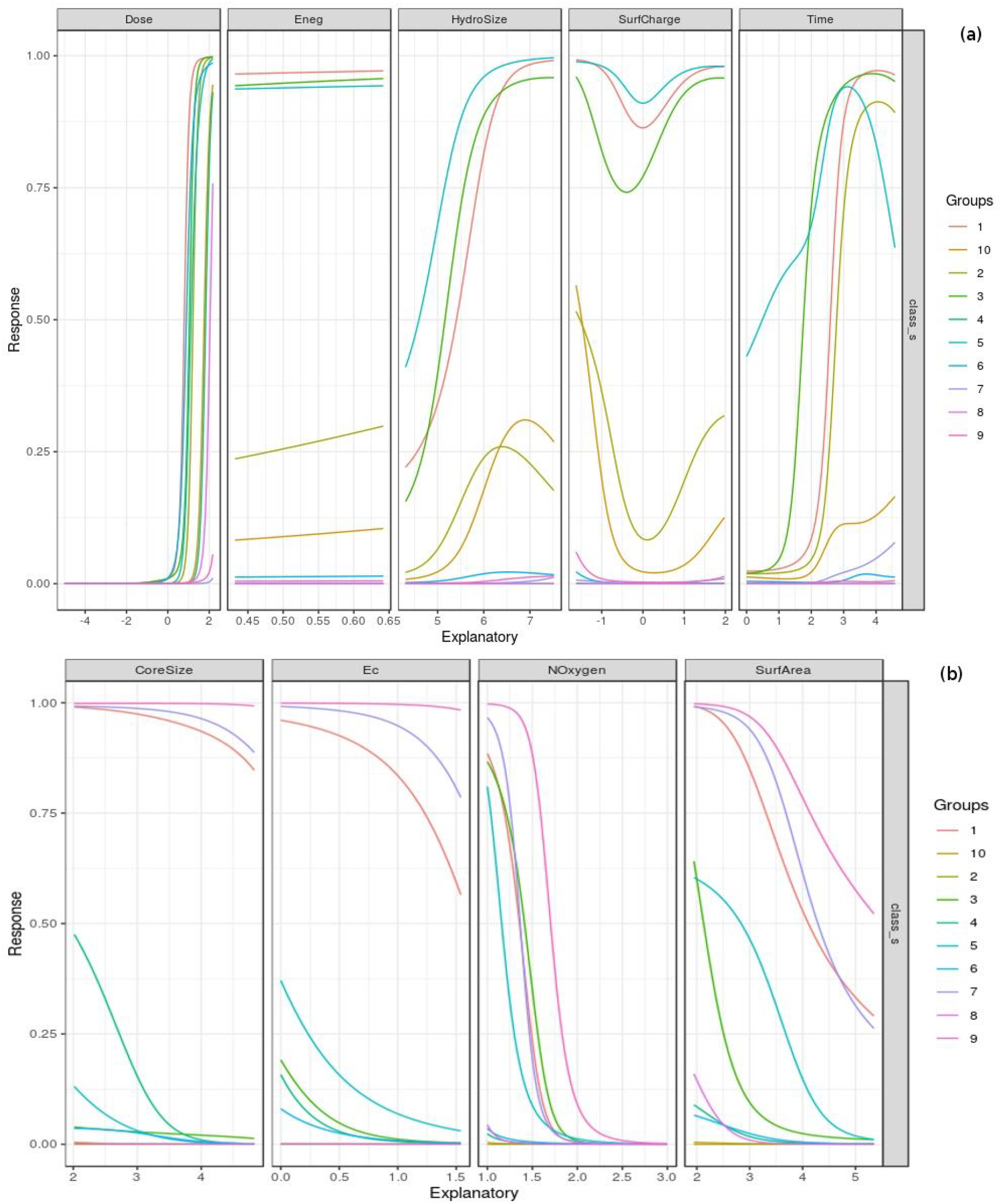
(a) Lek sensitivity analysis of attributes with positive effect on the outcome class. The steep effect of Dose is evident, with the location of the tipping point moving slightly with the cluster of the unevaluated variables. Increasing exposure times and HydroSize are also seen to tip to toxicity. (b) Lek sensitivity analysis of attributes with relatively consistent negative effect on the outcome class: CoreSize, Ec, NOxygen, and SurfArea. The number of lakes of the unevaluated variables is set to 10 in both the cases.

The two input variables that decisively differentiate the outcome are Dose and Noxygen. Dose appears to exert a nearly thresholding effect on the toxic class. The consistent sigmoidal effect seen in the ‘dose-response’ curve, independent of the lake of unevaluated explanatory variables, echoes the maxim attributed to Paracelsus, ‘The dose makes the poison.’ The attributes influencing toxicity also included: (i) Time, with a pronounced effect depending on the lakes of the unevaluated variables; and (ii) HydroSize, with a steady non-linear effect on toxicity that is also sensitive to the context of the unevaluated explanatory variables. The response profile for Eneg is almost flat at all lakes, indicating little to no effect in changing the outcome. The interpretation of the response with respect to SurfCharge remained obscure. NOxygen emerged as the attribute with the clearest inverse effect on toxicity, with a response profile displaying a tipping point to non-toxic class at most, but not all, of the centroids. Other attributes seen to dial down the toxicity include SurfArea, CoreSize, and Ec. These observations of effect size may be tempered with a significance analysis for deeper understanding.

In summary, the ML models of our work are represented by a purely numeric feature space of just nine predictors, and it is possible to consider them in their entirety, similar to the interpretability of a QSAR model. The models conform to the Findable, Accessible, Interoperable, Reusable (FAIR) principles, and are presented in a unified ensemble prediction engine, NanoTox (https://github.com/NanoTox). The scripts necessary for normalization and reproducible research are also available at NanoTox. Our methods may be extendable to other classes of engineered nanomaterials requiring urgent, sustainable, and rapid hazard classification prior to regular use [64–67].

## CONCLUSION

We have optimized the problem formulation of cytotoxicity modelling of nanoparticles using a principled approach agnostic of *in vitro* characteristics. The feature space is trimmed for multicollinearity, tunable hyperparameters were optimized, and the training data corrected for class imbalance. These steps further improved the performance of the parsimonious ML models to >96% balanced accuracy. The benefits of a parsimonious approach to modelling nanoparticle toxicity include enhanced model interpretability and generalizability. We have embedded our models into an unambiguous ensemble classifier that surpasses ~98% balanced accuracy. Our entire workflow is available as a free open-source resource for use and enhancement by the scientific community towards proactive non-invasive testing and design of nanoparticles for varied applications.

## Acknowledgements

We would like to thank the School of Chemical and BioTechnology, SASTRA Deemed University for infrastructure and computing support. This work was supported in part by the SASTRA TRR grant to A.P.

## Author contributions

A.P. conceived, designed and supervised the work. N.A. and A.P. performed research, and analyzed the results; A.P. wrote the paper.

